# Dietary restriction is evolutionary conserved on the phenotypic and mechanistic level

**DOI:** 10.1101/2025.03.26.645451

**Authors:** Sarah L Gautrey, Luke T Dunning, Toni I Gossmann, Mirre J P Simons

## Abstract

The anti-ageing response of Dietary Restriction (DR) is thought to be an ancient mechanistic response, reasoning from its phenotypic conservation in a wide range of organisms. However, DR is implemented using different diets and methods across species, and evidence for conservation at the mechanistic level remains limited. Here we tested the longevity and fecundity response to DR across eight different species of *Drosophila* using the same diets in a reaction norm framework. We confirm that DR is phenotypically conserved across *Drosophila*. Next, we used comparative transcriptomics across six species and found strongly concordant differential expression in response to DR (*r*_s_ > 0.28, < 0.72, P < 0.0001). We studied the evolutionary history of the top concordantly differentially expressed orthologous genes and identified that the large majority of these genes are “young” genes and are *Diptera* specific. Our results indicate that large parts of the DR response are likely to be taxonomic specific, suggesting that the genetic basis of DR is not widely conserved. To validate this hypothesis we tested whether the 15 most conserved genes that change in transcription in response to DR using conditional in vivo RNAi. Surprisingly, we found that 12 out of 15 genes tested had a lifespan phenotype, with 9 extending lifespan. Five of these genes are related to cysteine metabolism implicating it in the mechanisms of DR, further suggesting physiological compensation to DR is ubiquitous and providing a possible biomedical target. Our findings suggest that while large parts of the DR response are taxonomically specific, some core mechanisms are conserved across divergent species. The comparative approaches we used here hold promise to identify shared mechanisms relevant to our own species and therefore ultimately anti-ageing interventions.

## INTRODUCTION

Dietary restriction (DR), the reduction in food intake without malnutrition, has been shown to reliably extend health- and lifespan across a wide range of organisms, from single-celled yeast, to invertebrates, to mammals. As the DR response is found across widely divergent species its mechanisms are thought to be ancient and shared, and therefore translatable to human ageing research. The translational value of the mechanistic study of DR in other organisms depends heavily on the conservation of the underlying mechanism to our own species (*1*). The observed DR longevity response may have evolved once, in which case mechanisms might be conserved throughout species, alternatively it may be a case of recurrent convergent evolution (*2*).

When we understand the nutritional components and the molecular mechanisms responsible for DR we can start to develop drugs that mimic the health benefits of DR (*3*). Work on rodents set a widely held view that these life-extending effects were a result of overall caloric restriction, thus suggesting mechanisms underlying this extension involved protection from energy metabolism associated damage (*4*). However, since reduction of dietary yeast alone was shown to increase lifespan of flies, research has aimed to identify exactly which aspect of the restricted diet is important (*5, 6*). Experiments varying the ratio and amount of macronutrients in the diet have now shown that caloric intake does not necessarily determine the pro-longevity effect of DR and suggest that intake of the macronutrient protein is responsible (*5, 7–9*), and in particular essential amino acids (*10*), but see (*11*). However, protein restriction and caloric restriction (CR) are unlikely completely overlapping mechanisms in mammals (*12*). The effects of CR are larger than those achieved in macronutrient restriction (*7, 13*). When caloric restriction is imposed by lowering ambient temperature but feeding the exact same macronutrient composition a lifespan increase is observed (*14*).

Research into the mechanisms of DR has progressed in model organisms. For example: the DR response in *C. elegans* can be blocked by modulating alternative splicing via Splicing Factor 1 (*15*); *FGF21* is probably required for lifespan extension by DR in mice (*16*); Neuropeptide F signalling modulates DR in flies (*17*). Even though different species show similar responses on a physiological level (*18*), no single pharmaceutical or genetic manipulation negates or mimics the DR response across model organisms (*3, 19*). Perhaps the best example of this is a robust reduction in mTOR signalling that is seen when DR is imposed, but when manipulated DR and mTOR signalling independently affect longevity (*20–22*). As such there is a possible concern that identified mechanisms could be restricted or more dominant in certain species. Answering the question whether DR is conserved on a phenotypic and also mechanistic level is therefore highly relevant to understanding the biology of ageing.

The strongest effects of DR are found in the five most commonly studied model species, which questions the taxonomic generality of DR (*23, 24*). Thus, potentially the DR response is a laboratory artefact, perhaps resulting from relief of detrimental effects of overfeeding or other aspects of the lab environment (28, 29). However, DR has been shown to extend lifespan in these ‘model species’ even when collected from the wild, namely flies and C. elegans (*25, 26*), however DR did not extend lifespan of wild-derived killifish (*27*) and mice (*28*). Perhaps laboratory environments may simply make the effects of DR more pronounced, or diets are optimised (both rich and restricted) within each model species, strain, and maybe each specific laboratory to elicit the maximal lifespan extension (*29*). Indeed, in comparative studies in rotifers, multiple species did not show any increase in longevity, and there were high levels of intraspecific variation (*30, 31*). In general, substantial genetic variation is observed within species in response to DR, in both mice (*32*) and flies (*29, 33*), which is not expected when the response is highly conserved and under strong purifying selection.

A difficulty with interpreting both between-species and within-species variation in response to DR is the use of different diets and modes of restriction between species, and the use of often only two doses of diet (fully fed versus DR). DR shows a non-linear relationship with longevity and is commonly maximised at a relatively low concentration of dietary nutrients, with deviations on either side producing a decrease in lifespan either due to malnutrition or moving towards maximum reproduction or even overfeeding (*29, 34, 35*). Genotypes may differ in the position of maximum lifespan in nutritional space, with the full dose-response remaining intact (*29, 34, 35*). Therefore to appreciate if some genotypes or species do not respond to DR a range of diets and similar diet delivery is required (*29*). Here, we used multiple *Drosophila* species that can be kept on standard laboratory media and tested for conservation of the DR response on the phenotypic level in both lifespan and reproduction, as well as on the mechanistic level using transcriptomics and subsequent manipulation of identified mechanisms using functional genetics in flies (*Drosophila melanogaster*).

## RESULTS & DISCUSSION

### Phenotypic response to DR across species

Here we exposed eight different species of *Drosophila* to five different yeast concentrations in their diet to study the DR response. For five species we detected significant DR longevity responses (Figure 1). Two species (*D. pseudoobscure and D. yakuba*) showed minimal responses to diet. *D. mercatorum* lived longest on the highest yeast concentrations tested, suggesting a higher overall nutritional need (Figure 1, Table S1). For the species that showed a DR response, most had the strongest longevity response around 4% yeast, except for *D. willistoni* showing the strongest response on 2% yeast. These data fit with the hypothesis that reaction norms (dose-responses) to diet are critical when testing for the presence or absence of DR in certain contexts, genotypes or species (*29*). To test if species showed a true DR response (*35*), not a toxicity response to diet, we tested their egg laying response. All species on DR (diet of maximum longevity or 2% yeast if no clear response) showed a reduction in egg laying (Figure 2).

**Figure 1.**
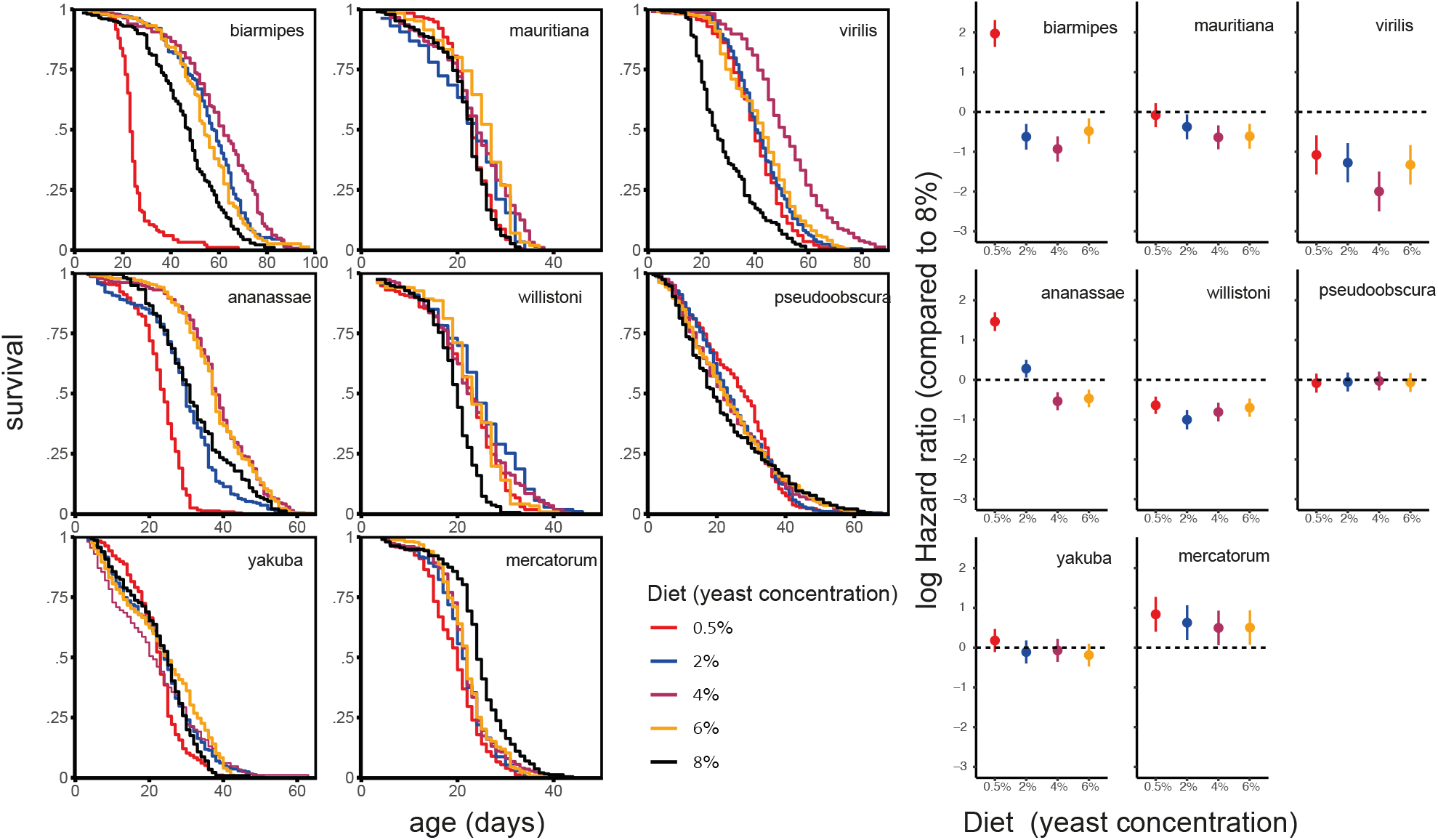
Survival curves (left set of panels) across *Drosophila species*, showing clear DR longevity responses in five of the eight species tested following classic reaction norms (*29, 34*). In model outputs from mixed-effects cox models (right) these effects can be most easily compared. Hazard ratios compared to the richest diet (8%) tested are given with their 95% confidence intervals (Table S1).

**Figure 2.**
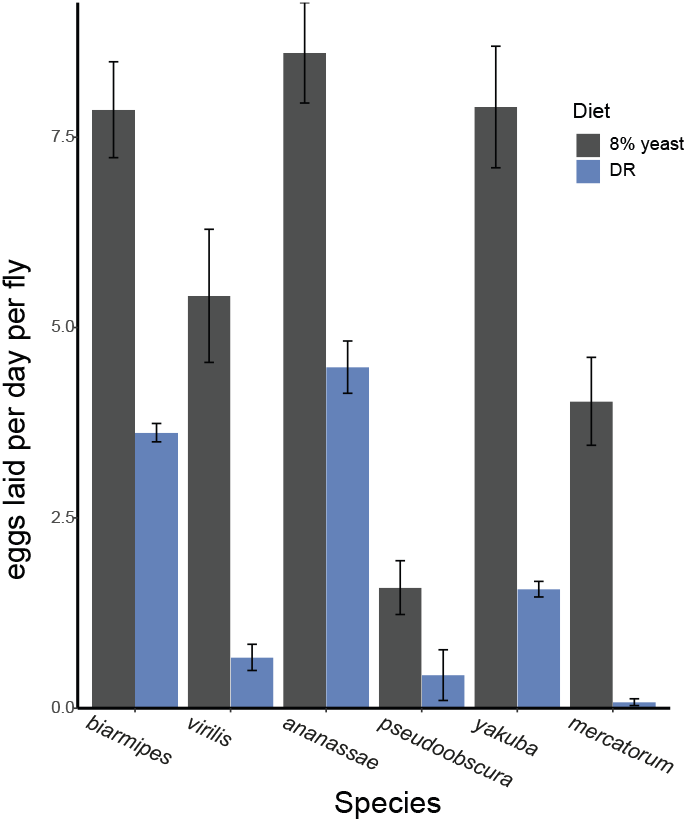
All species showed a reduction in egg laying in response to DR. This was statistically significant for each species (P < 0.05, Welch t-test) apart from *pseudoobscura*, which was not fecund. Inbreeding depression could explain low fecundity of this species and might also be the reason why this species was short-lived and unresponsive to DR. The same may hold for *yakuba*, even though did species was less fecund at DR. The strong reduction in egg laying seen from *mercatorum* does suggest that the rich diet was likely still not near their nutritional optimum (*35*).

### Transcriptomic response to DR

For five species that showed a clear response to DR we measured the transcriptomics response to DR and supplemented this dataset with *D. melanogaster* already available in the lab using the ywR strain that shows a consistent DR response (*36*). Orthologous genes across species were determined using protein sequence BLAST. Differential expression in response to DR was strongly positively correlated across species, suggesting shared mechanisms (Figure 3). To analyse which molecular pathways were affected across species we analysed the first principal component using gene set enrichment analysis (*37*) of KEGG (*38*) (Figure 4). Possibly in response to a change in reproductive output, homologous recombination and DNA replication are suppressed under DR (*39*). Transcription and translation in general appeared reduced, with suppression of ribosome and RNA related pathways (including the spliceosome). This is in contrast to prior work in humans and rhesus monkeys in which these pathways are upregulated (*40, 41*). However the overall suppression of protein synthesis through translation does fit with pro-longevity phenotypes attributed to DR (*18*), perhaps specifically through tRNA abundance (*42*). Autophagy genes were down in contrast with prior research, e.g. (*43*). Reduced expression of the foxO signalling pathway fits with reduced insulin/IGF signalling in response to, but not required for, DR (*44*). Upregulation of oxidative phosphorylation may fit with increased oxidative metabolism observed in caloric restriction and in long-lived mutant mice (*45*). Changes in expression are however difficult to directly relate to a change in the direction of affected physiology. As we argued previously using the example that an upregulation of DNA repair genes can either indicate a response to increased damage or a prioritisation of repair resulting in a healthier physiological state (*46*). Nevertheless, the enrichment of pathways previously associated with the lifespan benefits of DR suggests the transcriptomes measure the mechanistic response of DR successfully across species.

**Figure 3.**
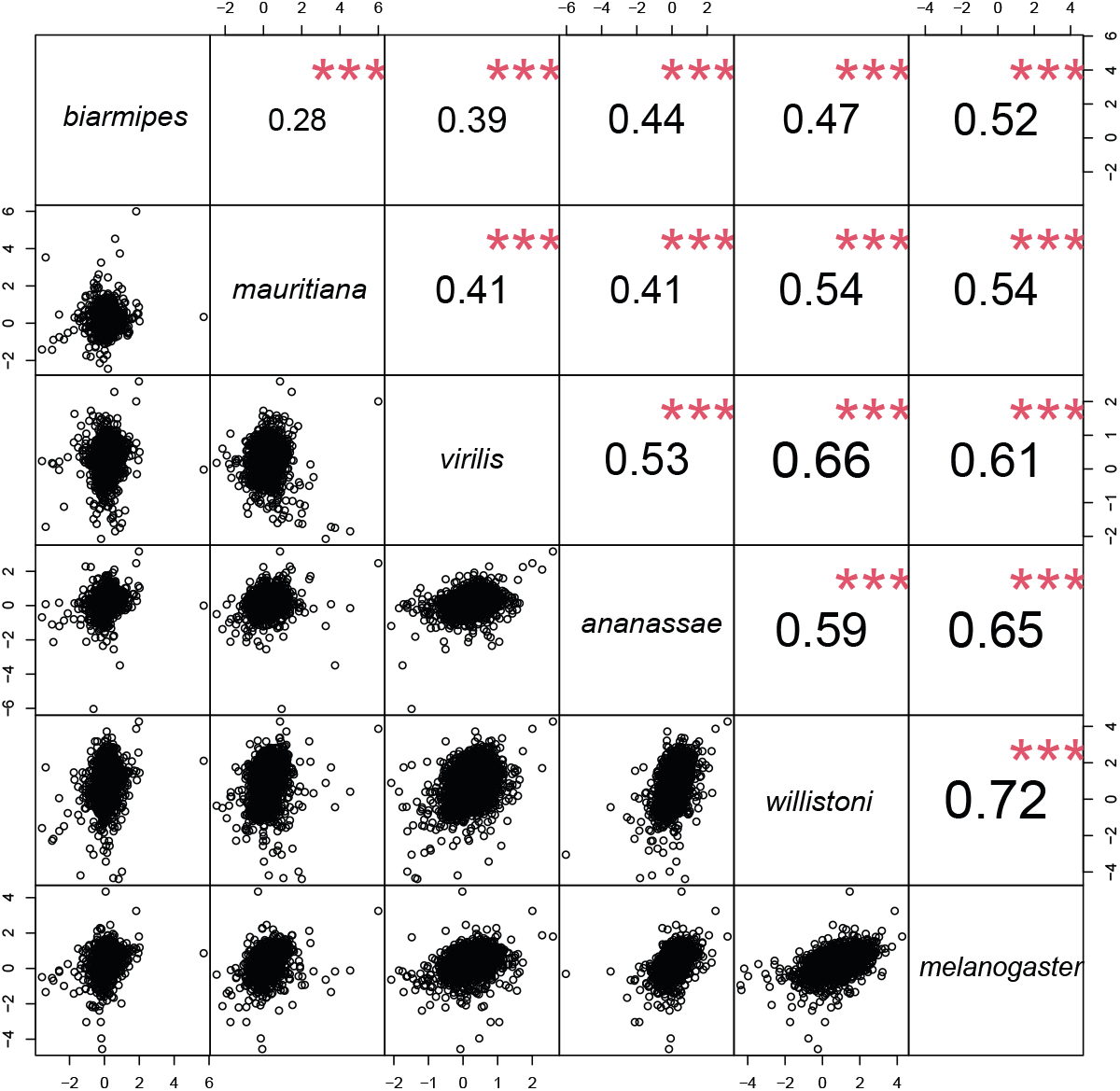
Rank correlations between differential expression in response to DR across species. Correlations were moderate to strong and highly significant for all comparisons. More closely related species did not show more strongly correlated responses (Figure S1).

**Figure 4.**
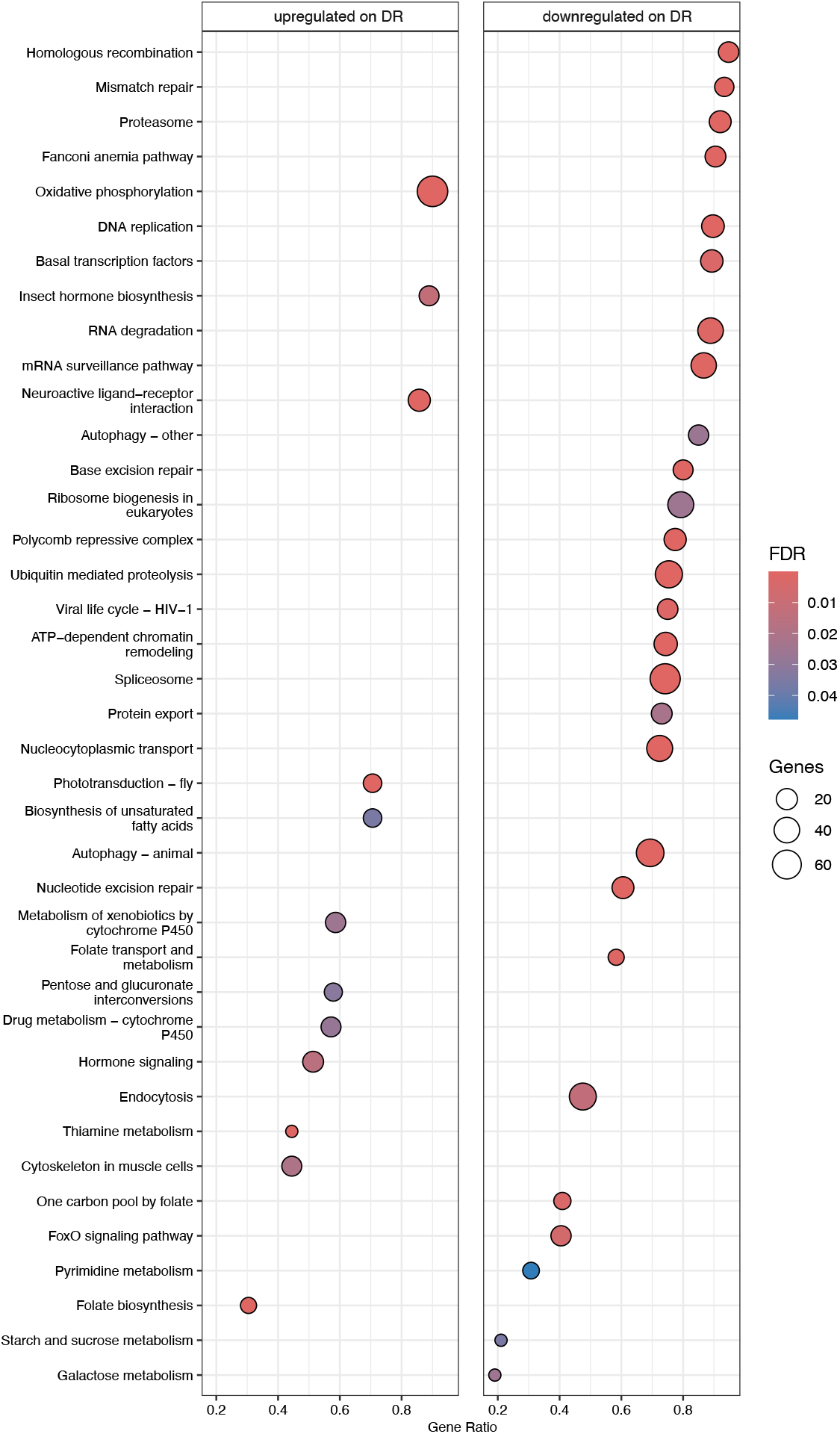
KEGG Gene Set Enrichment analysis for across the first principal component of differential expression in response to DR across species which describes the shared transcriptional change observed (Figure 3). Data is split for upregulated and downregulated genes. More traditional enrichment analysis dichotomising data between responsive and non-responsive genes disregarding direction of change gave similar results (Table S3).

### Evolutionary history and population genomics of genes responsive to DR

We analysed the evolutionary conservation of the genes that were found to be most responsive to DR. We selected the top 2.5% down and up regulated genes (total 5%) from the first principal component to dichotomise the data into responsive and non-responsive genes. Genes that changed in expression in response to DR were on average from younger phylostrata (*47*) (Figure 5A) meaning they were less conserved and have more recently evolved, or in other words not shared across all of life and more likely to be exclusive to arthropods. Alternatively, using a continuous scale instead of dichotomising, the rank correlation of differential expression between species was stronger for the younger phylostrata, peaking for genes that are unique to Arthropods only (Figure 5B). These findings further fit with results we obtained from a more granulated analysis we did using the classic comparative genomic prior analysis across twelve *Drosophila* species (*48*). The amount of non-synonymous over synonymous mutations (dN/dS) was higher in the DR responsive genes (as determined by PCA; Wilcoxon rank test, p < 0.0001, Figure 5C), and this was driven by a change in dN (P < 0.0001) and not dS (P = 0.64). This indicates a relaxation in purifying selection across species or possibly a small shift towards more adaptive evolution, both fitting with the idea that the genes underlying the mechanisms of DR are relatively young and not strongly conserved. Similar results were found on a population level using the rich *Drosophila melanogaster* genomic resource the DPGP3 (*49*) with transcripts that responded significantly to DR in *melanogaster* showing a small but significant increase in both Tajima’s D and P_N_ / P_s_ (Figure 5D, E), fitting with relaxed purifying selection.

**Figure 5.**
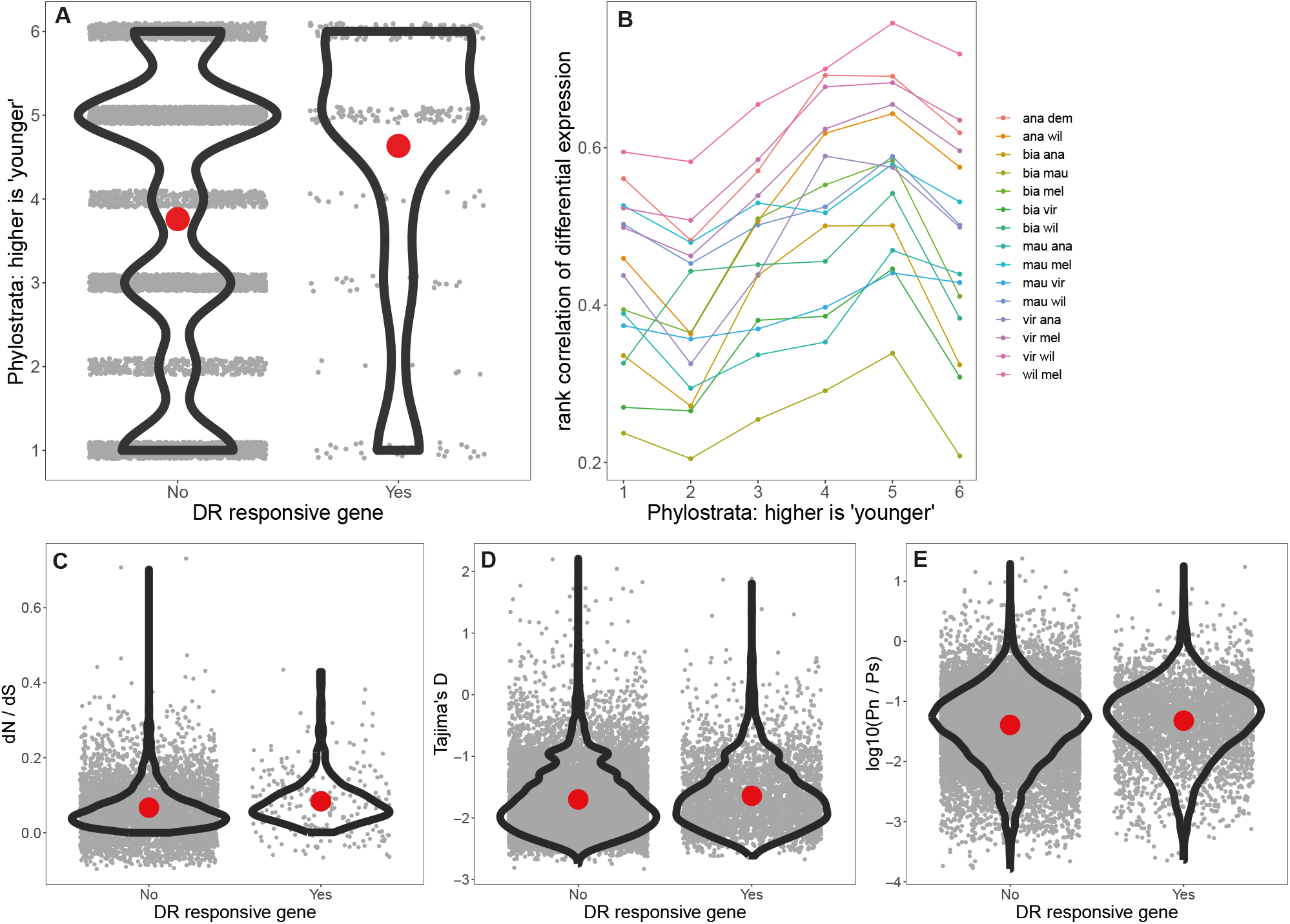
**A**. Genes responsive to DR as determined by the first principal component belonged to higher phylostrata, meaning younger, less conserved genes (Wilcoxon rank test, P < 0.0001). **B**. Genes belonging to each phylostrata showed different levels of concordance between species in total differential expression in response to DR (Kruskall-Wallis; chi-squared = 17.8, df = 5, P < 0.005). Each individual strata differed significantly from the next (pair-wise Wilcoxon rank exact test, p < 0.001) peaking at phylostrata five which is Arthropod specific. Phylostrata are based on published work (*47*): 1) Common ancestor to Fungi, Plants & Eumetazoa, 2) excluding Plants, 3) excluding Fungi, 4) Bilateria, 5) Arthropoda, 6) Dipetera. **C**. dN/dS determined across twelve Drosophila species is higher for DR responsive genes **D**. Within Drosophila melanogaster using simple significance cut-offs to determine DR responsive genes, both Tajima’s D and P_N_ / P_s_ were higher (**D & E**).

### Testing for possible conserved mechanisms of dietary restriction

On average the genes that change in transcription in response to DR across species are relatively young and less conserved due to relaxed purifying selection. It is possible however, that species, or taxon-specific signals mask shared conserved mechanisms. We therefore identified a top set of 15 genes that responded to DR and were widely conserved across divergent species (belonged to the “oldest” phylostrata) using a combination of principal component and significance testing (see methods). As a test that these genes would represent the “ancient” highly conserved molecular machinery required for DR we manipulated each of these genes using functional genetics under both DR and fully fed conditions in *Drosophila melanogaster*. In addition to this evolutionary prediction, any novel mechanism identified that is involved in the DR response of these conserved genes will be of strong biomedical interest, as the mechanisms likely apply to our own species. Twelve out of the fifteen genes tested had a longevity phenotype, with eight extending lifespan on DR (Figure 6). Three genes showed no phenotype, and three lived shorter. These results indicate that at least part of the conserved machinery responsive to DR modulates lifespan.

**Figure 6.**
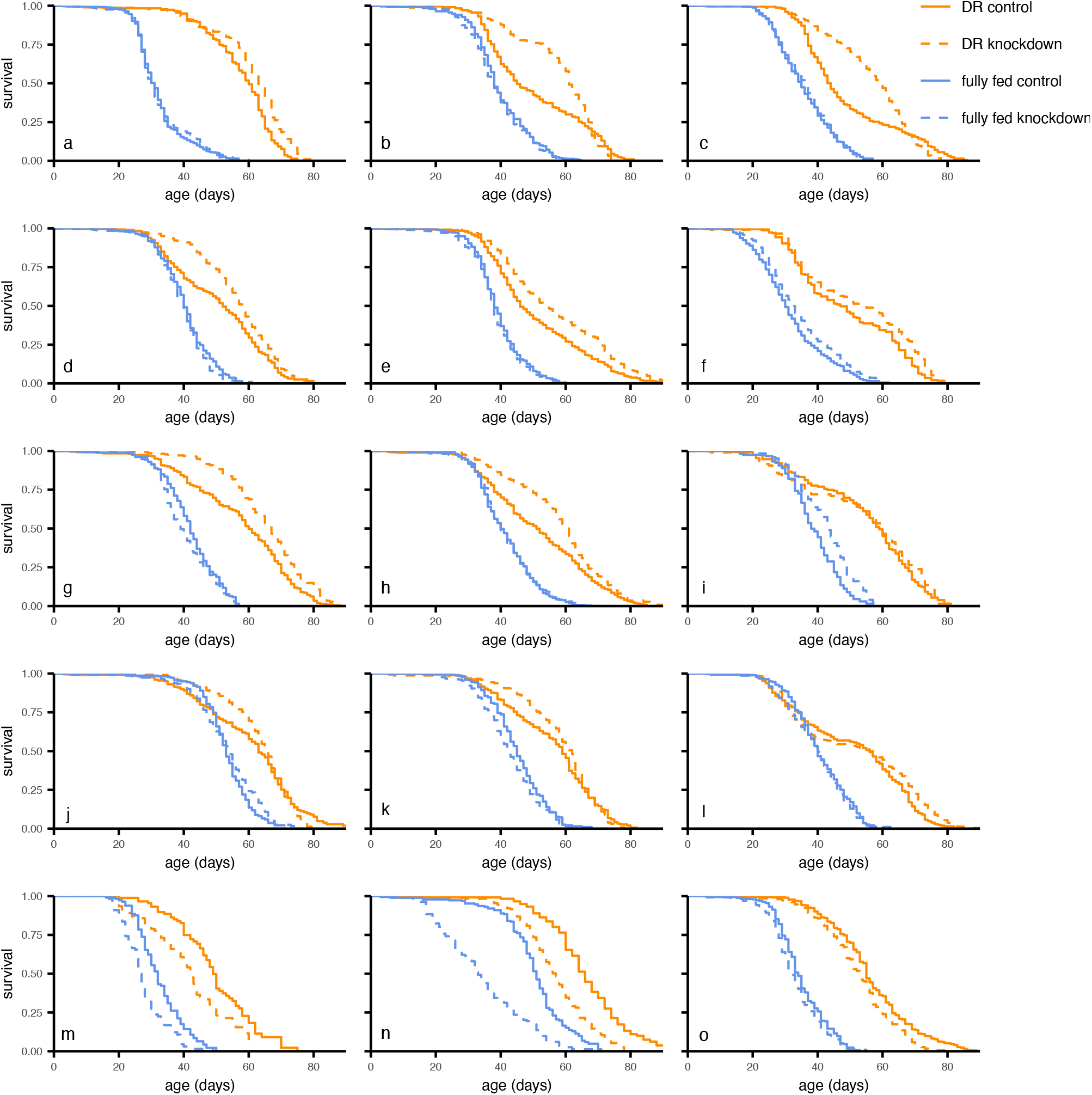
Test of candidate genes using conditional (GeneSwitch) knockdown (*in vivo* RNAi). Many of the genes tested showed lifespan phenotypes, with the most common an extension of lifespan on DR. Statistics provided in Table S2.

### Core mechanisms of DR

One of the most striking findings is that a large proportion of the genes tested extended lifespan on DR when knocked down (9 out of 15 genes, Figure 6a-i). Interestingly, five of these genes respond to DR by increasing in transcription (Figure 6a,c,f,h,i; Table S2). Thus, in effect we are suppressing the natural transcriptomic response to DR by RNAi. These results can be interpreted as reducing compensatory responses to DR, therefore allowing the DR longevity phenotype to become more prominent. These effects are interesting, especially for translational purposes, but perhaps should not be considered core mechanisms of DR. In contrast, genes that do go down in expression under DR, and then extend longevity when knocked down could be part of the core machinery of DR (Figure 6b,d,e,g).

Still the classic interpretation of such core machinery of DR would also mean that under fully fed conditions, flies would be pushed into a DR-like state by knockdown of these candidate genes. The knockdown of only two genes extended lifespan on rich diets (Figure 6f,i) and for these two genes, this classic interpretation does not hold. Their transcription goes up in response to DR, and hence a reduction in expression through knockdown should push flies away from a DR-like state. Again, perhaps DR is activating compensatory mechanisms to resist the DR-like state, explaining why a suppression of such mechanisms would lead to a lifespan extension. Three genes (Figure 6m,n,o) appeared essential and knockdown led to a truncation of lifespan on both diets.

The exact dose and direction of manipulation (using overexpression) could be crucial for the effects of these individual genes. Further research will be required to delineate the exact mechanisms. It is striking however that amino-acid metabolism (including proteolysis) and carbohydrate metabolism is a common theme amongst our validated candidate genes (Table S2). The direction of transcriptional change of these genes is not unidirectional, perhaps explaining why in directional analysis of KEGG pathways these terms are not prominent (Figure 4). Carbohydrate, amino-acids and specific nutrient metabolism are slightly more enriched using a non-directional KEGG enrichment of the first principal component across species (Table S3). Perhaps these changes represent a shift towards aspects of nutrient metabolism, or they represent differential compensatory and causal mechanisms. Whether amino-acid metabolism or other nutrients are central to DR remains to be fully determined (*11, 18, 29, 50–52*).

From our functional genetic tests of candidate genes identified as DR responsive and conserved it appears they are enriched for longevity phenotypes. However, we did not formally test for this as we did not take 15 random genes and also tested them similarly. Still, our mechanistic tests do support the notion that the mechanisms of DR could be widely conserved. The transcriptional signature of DR across species appears to include many species- or taxon-specific signals that mask shared conserved mechanisms. DR-responsive genes that are conserved appear to hold important mechanistic insights into the biology of ageing.

A prominent enrichment in the candidate gene set and KEGG analysis (Figure 4, Table S3), is metabolism related to cysteine. The orthologs of *Sobremesa* and *Maltase A6* in humans (*SLC7A9* and *SLC3A1*) are involved in familial cystinuria (the deposit of cystine, the oxidised form of cysteine, in renal tubules) (*53*). Both *SLC7A9* and *SLC3A1* form an active transport complex together that regulates transport of amino-acids in the intestine and the kidney driven by intracellular amino-acid concentrations and the reduction of cystine to cysteine. The ortholog of *CG5493* is *CDO1* (Cysteine dioxygenase) metabolising cysteine to cysteine sulfinic acid which can be converted to taurine or sulphate and pyruvate (*54*). *MTHFD1* is the ortholog of pug, an enzyme in folate metabolism, which links to cysteine via the methionine cycle. Interestingly, *MTHFD1* loss of function mutant mice have higher levels of cysteine, but lower levels of methionine (*55*).

*CG10621* probably produces an homocysteine S-methyltransferase enzyme, converting cysteine to methionine. *CG10621* expression responds to methionine restriction (*49*, reduced) and tyrosine restriction (*55*, increased) in opposite directions. Furthermore, serine is required to produce cysteine (*57*), and *astray* codes for the final enzymatic step to produce serine from carbohydrates.

Each of the enzymes mentioned above reduced in expression in response to DR, and both transporter genes responded in opposite directions (Table S2). Consistently across these enzymes further reduction of expression using knockdown led to a further extension of lifespan on DR. Apart from *astray* a reduction in each of these enzymes (*CG5493, pug, CG10621*) is predicted to lead to higher internal levels of cysteine. The reduction in *astray* is possibly a response to carbohydrate levels and not a response to cysteine-associated metabolism. Therefore, we hypothesise that the combined mechanism we distilled here is a response to defend cysteine levels in the face of DR. The reduction in specific metabolic conversions of cysteine, namely the production of taurine (*58*) (*CG5493*) and alterations in the methionine cycle (*59, 60*) (*pug* and *CG10621*), may be responsible for the longevity benefit of DR. Alternatively, different metabolic uses of cysteine that may receive more metabolic flux, namely hydrogen sulfide (*61*) and glutathione (*62*) have been associated with healthy ageing.

## Conclusions

We show here for the first time that DR is phenotypically and mechanistically conserved using *Drosophila*, a model species in the biology of aging, of which conveniently widely divergent species can be fed the exact same diet. Surprisingly, the genes that respond transcriptionally to DR are not strongly conserved. Yet, when we tested highly conserved genes only using functional genetics in *melanogaster* we find that these identified candidate genes modulate lifespan. However, our findings do not support the idea of a master regulator for DR (*19*), as experimental reduction in expression of the majority of genes, which naturally are upregulated in response to DR, enhances rather than suppresses DR’s longevity benefits. We interpret this as limiting the compensatory physiology naturally activated under DR, in effect removing restrictions on how DR exerts its longevity benefits. In general, it is probable that organisms have evolved to actively resist a ‘healthier’ state in order to avoid reproductive fitness costs (*63*). They defend their physiology through compensatory mechanisms, resisting the longevity benefits of DR. The restriction of specific nutrients will likely lead to an upregulation of transport and recycling of scarce nutrients. We have previously hypothesised that such mechanisms could be responsible for the costs of the refeeding syndrome observed when organisms are moved from DR to fully fed conditions (*46*).

Similar compensatory mechanisms also explain why a large part of the DR response is taxon-specific, as some of this compensation is either more important for fitness in some taxa or some physiology (e.g. egg production) may not even exist in other taxa. From a translational perspective the reason why DR needs to be substantially restrictive in nutritional space could be that it has to overcome internal compensatory mechanisms before longevity benefits are reached. Compensatory mechanisms that limit DR could therefore be exceptional targets to achieve the health benefits of DR without the need for highly restrictive diets. Discrepancies in how individual nutrients modulate the DR response (*10, 11*) may also be explained by how and in which part of the nutritional space compensatory mechanisms become exhausted or costly (*29*). For the evolution of DR, our work supports the idea of conserved mechanisms, whilst at the same time pointing to taxon and species-specific, as well as compensatory physiology, masking this shared physiology. Unmasking these effects using the approach we took here holds promise to identify compensatory mechanisms in response to DR that are shared between model species and our own species, to eventually contribute to anti-ageing interventions.

## METHODS

### Fly husbandry, diet and environment

Eight species of the *Drosophila* genus were obtained from Cornell Drosophila Stock Center(*48*). All species were cultured on rich yeast media (8% autolysed yeast, 13% table sugar, 6% cornmeal, 1% agar and niapagin 0.225%) (*35, 46*). Cooked fly media was stored for up to two weeks at 4-6°C, and warmed to 25°C before use. Experimental diets had the same composition, but yeast was varied, 0.5%, 2%. 4%, 6% or 8%, as this is shown to be the main determinant of the DR longevity response in flies (*35, 64*). Experiments were conducted in a climate-controlled environment with a 12:12 hour light-dark cycle, temperature at 25°C and 50-60% relative humidity.

### Across species longevity in response to DR

All species of *Drosophila* were grown in bottles containing a rich diet (8% yeast), and kept at 25°C. In each of these bottles, 15 females and 5 males were allowed to lay eggs for 4 days. When offspring began to eclose, individuals were transferred to mating bottles, where they were left to mate for ∼48 hours. This was repeated every day to make age-matched cohorts. Flies were then sorted under carbon dioxide anaesthesia (Flystuff Flowbuddy; <5L / min), and females were transferred to demography cages, specially designed to allow removal of deceased flies, and changing of food vial with minimal disturbance to living flies (*65*). Up to 100 female flies were housed in each cage, with up to 4 cages per treatment, depending on success of fly population growth. Experimental diet (0.5%, 2%, 4%, 6% or 8% yeast) began 1-2 days after flies were transferred into cages. At the same time, any dead flies were removed from the cages, recorded and discarded. This was repeated on alternate days, with the food vial changed each time. Living flies which were stuck to the food vial or died as a result of sticking to the food or becoming trapped in any part of the cage, and flies which escaped were right-censored (*46*).

### Across species fecundity in response to DR

Following the longevity experiment we used six of the eight *Drosophila* species: *ananassae, yakuba, biarmipes, virilis, mercatorum and pseudoobscura*; 3 of which demonstrated increased longevity with DR, and 3 which did not. This experiment had the same initial set up, in the way flies were grown and sorted. Flies were then put in demography cages with ∼25 flies per cage, and 3 cages per treatment. Each cage was allocated one of two diets; fully fed conditions (8% yeast), and DR (the yeast concentration that resulted in the longest lifespan extension, or 2% yeast if no effect was seen). Individuals were left to acclimatise to the diet for 6 days, with fresh food being given on alternate days, and deceased individuals removed and recorded similarly to the previous experiment. From day 6 the number of eggs were counted every 24h until Day 10, using a dissection microscope, and the egg count was averaged per cage across days for analysis.

### Data Analysis of survival and egg laying

All analyses and data handling were carried out using R. Survival associated with varying levels of dietary restriction was analysed using mixed cox-proportional hazard models (*66*), which used ‘cage’ as a random term to avoid pseudo-replicated effects. Date of sorting into cages was fitted as a fixed effect in the models which further corrects for shared environmental effects. Egg laying was analysed using a Welch’s t-tests, which allowed comparison of the mean number of eggs laid per day between two dietary regimes.

### Comparative transcriptomics of DR

Flies were sampled from their cages between age 20 and 27 days, with DR and fully fed diet sampled being age paired-matched within each species. Samples were processed as a mix of 4 to 5 individuals, snap frozen and kept until analysis at -80C. We analysed six species and used the diet optimal for DR compared to 8% yeast for all. Sample size and stock information is provided in Table S4. RNA was extracted using Qiagen RNeasy mini kits. RNA quality was validated and concentration was determined using Agilent Tapestation. Library prep and sequencing was carried out by the Oxford Genomics Centre at the Wellcome Centre for Human Genetics on the Illumina Novaseq platform using 150bp paired end. For bioinformatic analysis we used hisat2, stringtie and ballgown (*67*) and we used respective genomes and their associated transcript annotation files as reference files. Differential expression analysis was carried out using edgeR and ‘glmFit’ (*68*). We then analysed differential expression data using correlation analysis and principal component analysis using the rationale described in the results section.

### Test of candidate genes in Drosophila melanogaster

We selected a limited set of genes that were DR responsive (the top 2.5% down and up regulated genes (total 5%) from the first principal component) that were most conserved (in the first three phylostrata). We further selected for genes that both had a concordant sign of differential expression and were statistically significant (ɑ = 0.05) in five or more species and again were most conserved (in the first three phylostrata). Then finally we excluded genes annotated as involved in meiosis/mitosis as this is likely a signal related to the ovary only reflecting the change seen in reproduction under DR. As a pragmatic further consideration we tested only genes that had in vivo RNAi constructs available from the TRiP collection (*69*) ordered from the Bloomington Drosophila Stock Centre. GeneSwitch (GS) allowed us to conditionally activate the RNAi, by adding the drug RU486 (mifepristone) to the diet, thus preventing confounding effects of genetic background mutations (*70, 71*). We used da-GS, the daughterless (da) promoter is active in almost all cells of the fly and the GeneSwitch construct will thus be present in each cell of the fly allowing us to conditionally knockdown candidate genes using RNAi in the whole fly (*72*). We measured mortality on female flies in each of these genetic crosses, with 4 dietary conditions; 2% and 8% yeast, each with and without addition of RU486 (at 200µM final media concentration). Diets are composed of sugar, yeast, cornmeal, agar and nipagin (*11, 46*). Non-RU diets have addition of ethanol as a control, and were split from the same media batch. Prior work from others and more importantly our current lab has shown for other candidates that this system is highly reliable, verifiable using qPCR and RNAseq (unpublished and (*73*)). For the purposes of this study we assume that the target genes are manipulated as intended. RU486 itself does not affect fly longevity, and we confirmed this in our lab previously (*73, 74*), and here using a cross of da-GS with empty attp2 vector on both 2% and 8% yeast conditions (Figure S2). Flies were housed in cages designed to easily remove dead flies and change food with minimal disturbance to living flies, with up to 100 flies in each cage and 5 cages per treatment (20 per cross). Mortality was measured on alternate days until all flies in the cage had died.

## ACKNOWLEDGEMENTS

We thank Laura Hartshorne, Andrew McCracken, Alexander Charles, Gracie Adams and other members of Simons lab group for continued support. We thank the Oxford Genomics Centre at the Wellcome Centre for Human Genetics (funded by Wellcome Trust grant reference 203141/Z/16/Z) for the generation and initial processing of the sequencing data. We thank the Cornell Drosophila Stock Center and the Bloomington Stock Center.

## AUTHOR CONTRIBUTIONS

MJPS designed the study, conducted experiments, analysed the data and wrote the manuscript. SG conducted experiments, wrote a first draft of the manuscript and analysed data. TG and LD helped design the study and supported the comparative and evolutionary genetics and genomics.

