## Supplementary Material for "Dietary restriction is evolutionary conserved on the phenotypic and mechanistic level"

**Table S1**. Log hazard ratios from 'coxme' models testing the effect of lowering yeast concentration in the diet (DR) against fully fed conditions (8% yeast). Statistics correspond to Figure 1.

| | estimate | exp (estimate) | se | p |
|---|---|---|---|---|
| ***D. ananassae*** | | | | |
| 0.5% yeast | 1.462 | 4.316 | 0.120 | <0.001 |
| 2% yeast | 0.281 | 1.324 | 0.114 | <0.001 |
| 4% yeast | -0.538 | 0.584 | 0.113 | <0.001 |
| 6% yeast | -0.469 | 0.626 | 0.113 | <0.001 |
| ***D. yakuba*** | | | | |
| 0.5% yeast | 0.178 | 1.200 | 0.148 | 0.23 |
| 2% yeast | -0.111 | 0.895 | 0.148 | 0.45 |
| 4% yeast | -0.070 | 0.933 | 0.149 | 0.64 |
| 6% yeast | -0.188 | 0.828 | 0.146 | 0.20 |
| ***D. biarmipes*** | | | | |
| 0.5% yeast | 1.973 | 7.191 | 0.172 | <0.001 |
| 2% yeast | -0.622 | 0.537 | 0.164 | <0.001 |
| 4% yeast | -0.930 | 0.395 | 0.163 | <0.001 |
| 6% yeast | -0.478 | 0.620 | 0.162 | <0.001 |
| ***D. mercatorum*** | | | | |
| 0.5% yeast | 0.833 | 2.301 | 0.223 | <0.001 |
| 2% yeast | 0.624 | 1.866 | 0.223 | 0.0052 |
| 4% yeast | 0.493 | 1.637 | 0.221 | 0.026 |
| 6% yeast | 0.502 | 1.652 | 0.221 | 0.023 |
| ***D. pseudoobscura*** | | | | |
| 0.5% yeast | -0.084 | 0.920 | 0.122 | 0.49 |
| 2% yeast | -0.055 | 0.947 | 0.121 | 0.65 |
| 4% yeast | -0.031 | 0.970 | 0.121 | 0.80 |
| 6% yeast | -0.067 | 0.935 | 0.122 | 0.58 |
| ***D. virilis*** | | | | |
| 0.5% yeast | -1.080 | 0.340 | 0.251 | <0.001 |
| 2% yeast | -1.277 | 0.280 | 0.251 | <0.001 |
| 4% yeast | -2.000 | 0.136 | 0.255 | <0.001 |
| 6% yeast | -1.327 | 0.265 | 0.253 | <0.001 |
| ***D. mauritiana*** | | | | |
| 0.5% yeast | -0.080 | 0.924 | 0.154 | <0.001 |
| 2% yeast | -0.373 | 0.688 | 0.160 | <0.001 |
| 4% yeast | -0.637 | 0.529 | 0.153 | <0.001 |
| 6% yeast | -0.611 | 0.543 | 0.159 | <0.001 |
| ***D. willistoni*** | | | | |
| 0.5% yeast | -0.640 | 0.527 | 0.111 | <0.001 |
| 2% yeast | -1.001 | 0.367 | 0.125 | <0.001 |
| 4% yeast | -0.811 | 0.444 | 0.112 | <0.001 |
| 6% yeast | -0.701 | 0.496 | 0.117 | <0.001 |

**Table S2.** Set of most highly conserved, DR responsive genes from six species of the Drosophila lineage tested using conditional in vivo knockdown in melanogaster. Plot refers to the panel in Figure 5. The effects on longevity are reported for knockdown for each gene along with a summary from Flybase. The expression change is reported as logFC compared to a fully fed diet (8% yeast) for *melanogaster*.

| Plot | Bloomington Stock | Flybase gene ID | LogFC (DR effect in D. mel) | Name | Effect on DR (logHR) | Effect on fully-fed conditions ((logHR) | Gene Summary from Flybase |
|---|---|---|---|---|---|---|---|
| a | 25850 | FBgn0010015 | 1.120 | Calcineurin A1 | -0.48 ± 0.088, P < 0.0001 | -0.033 ± 0.078, P = 0.67 | Calcium-dependent, calmodulin-stimulated protein phosphatase. This subunit may have a role in the calmodulin activation of calcineurin. |
| b | 38338 | FBgn0023129 | -0.62 | astray | -0.25 ± 0.11, P = 0.021 | 0.087 ± 0.10, P = 0.85 | Catalyzes the last step in the biosynthesis of serine from carbohydrates. The reaction mechanism proceeds via the formation of a phosphoryl-enzyme intermediates |
| c | 41695 | FBgn0004117 | 1.192 | Tropomyosin 2 | -0.31 ± 0.10, P = 0.003 | -0.05 ± 0.10, P = 0.61 | Tropomyosin, in association with the troponin complex, plays a central role in the calcium dependent regulation of muscle contraction. May also regulate motor systems required to maintain nuclear integrity and apico-basal polarity during embryogenesis. |
| d | 42011 | FBgn0032726 | -1.600 | CG10621 | -0.31 ± 0.09, P = 0.001 | 0.079 ± 0.091, P = 0.39 | Predicted to enable S-adenosylmethionine-homocysteine S-methyltransferase activity. Predicted to be involved in S-methylmethionine cycle and methionine biosynthetic process. |
| e | 53291 | FBgn0020385 | -0.27 | pug | -0.36 ± 0.087, P < 0.001 | 0.040 ± 0.087, P = 0.64 | encodes the trifunctional enzyme methylenetetrahydrofolate dehydrogenase involved in the pigmentation of pteridines and ommochromes |
| f | 57404 | FBgn0001187 | 0.647 | Hexokinase C | -0.27 ± 0.12, P = 0.030 | -0.21 ± 0.19, P = 0.041 | Hexokinase C (Hex-C) encodes a hexokinase involved in glucose homeostasis. |
| g | 57735 | FBgn0034364 | -1.076 | CG5493 | -0.46 ± 0.14, P = 0.0014 | 0.30 ± 0.14, P = 0.034 | Predicted to enable cysteine dioxygenase activity and ferrous iron binding activity. Predicted to be involved in L-cysteine catabolic process. |
| h | 51791 | FBgn0030574 | 0.836 | Sobremesa | -0.36 ± 0.13, P = 0.0058 | -0.06 ± 0.13, P = 0.59 | Predicted to enable L-amino acid transmembrane transporter activity. Predicted to be involved in amino acid transmembrane transport. Located in cell cortex. |
| i | 60043 | FBgn0032144 | 1.326 | CG17633 | -0.30 ± 0.15, P = 0.047 | -0.43 ± 0.14, P = 0.0023 | Predicted to enable metallocarboxypeptidase activity. Predicted to be involved in proteolysis. Predicted to be active in extracellular space. |
| j | 55319 | FBgn0022160 | 0.650 | Glycerophosphate oxidase 1 | 0.02 ± 0.14, P = 0.85 | -0.19 ± 0.13, P = 0.15 | Glycerophosphate oxidase 1 (Gpo1) encodes a mitochondrial inner membrane protein with glycerol-3-phosphate dehydrogenase activity. |
| k | 55346 | FBgn0032381 | 0.243 | Maltase B1 | -0.066 ± 0.16, P = 0.68 | 0.23 ± 0.16, P = 0.13 | Predicted to enable maltose alpha-glucosidase activity. Predicted to be involved in carbohydrate metabolic process. |
| l | 57563 | FBgn0034438 | 0.715 | CG9416 | -0.21 ± 0.12, P = 0.086 | 0.081 ± 0.12, P = 0.50 | Predicted to enable metalloexopeptidase activity. Predicted to be involved in proteolysis. |
| m | 34740 | FBgn0039580 | -0.012 | Glutamine:fructose-6-phosphate aminotransferase 2 | 0.67 ± 0.23, P = 0.0034 | 0.88 ± 0.20, P < 0.001 | Enables glutamine-fructose-6-phosphate transaminase (isomerizing) activity. Predicted to be involved in UDP-N-acetylglucosamine metabolic process; fructose 6-phosphate metabolic process; and protein N-linked glycosylation. |
| n | 60398 | FBgn0050360 | -0.693 | Maltase A6 | 0.76 ± 0.31, P = 0.014 | 1.24 ± 0.27, P < 0.001 | Predicted to enable maltose alpha-glucosidase activity. Predicted to be involved in carbohydrate metabolic process. |
| o | 44495 | FBgn0034497 | 1.339 | Mitochondrial phosphate carrier protein 1 | 0.35 ± 0.08, P < 0.001 | 0.19 ± 0.089, P = 0.035 | Predicted to enable inorganic phosphate transmembrane transporter activity. Predicted to be involved in phosphate ion transmembrane transport. Predicted to be located in mitochondrion. Predicted to be active in mitochondrial inner membrane. |

**Table S3.** Enrichment of across species genes that respond transcriptionally to DR disregarding direction.

| KEGG Pathway | Universe | PC1 <2.5% >97.5% | P |
|---|---|---|---|
| DNA replication | 29 | 16 | <0.0001 |
| Base excision repair | 20 | 6 | <0.0001 |
| Mismatch repair | 15 | 5 | 0.001 |
| Thiamine metabolism | 9 | 4 | 0.001 |
| Galactose metabolism | 21 | 5 | 0.003 |
| One carbon pool by folate | 22 | 5 | 0.004 |
| Folate biosynthesis | 23 | 5 | 0.005 |
| Nucleotide excision repair | 38 | 6 | 0.011 |
| Homologous recombination | 19 | 4 | 0.013 |
| Starch and sucrose metabolism | 19 | 4 | 0.013 |
| Folate transport and metabolism | 12 | 3 | 0.020 |
| Cysteine and methionine metabolism | 26 | 4 | 0.039 |

**Table S4.** Detailed Information about the RNAseq experiment across *Drosophila* species.

| Species | Total sample size | DR diet | Stock |
|---|---|---|---|
| *ananassae* | 6 | 4% | 14024-0371.34 |
| *biarmipes* | 4 | 4% | 14023-0361.09 |
| *virilis* | 6 | 4% | 15010-1051.88 |
| *mauritiana* | 4 | 6% | 14021-0241.151 |
| *willistoni* | 4 | 2% | 14030-0811.24 |
| *melanogaster* | 20 | 2% | ywR (Phillips & Simons 2024) |

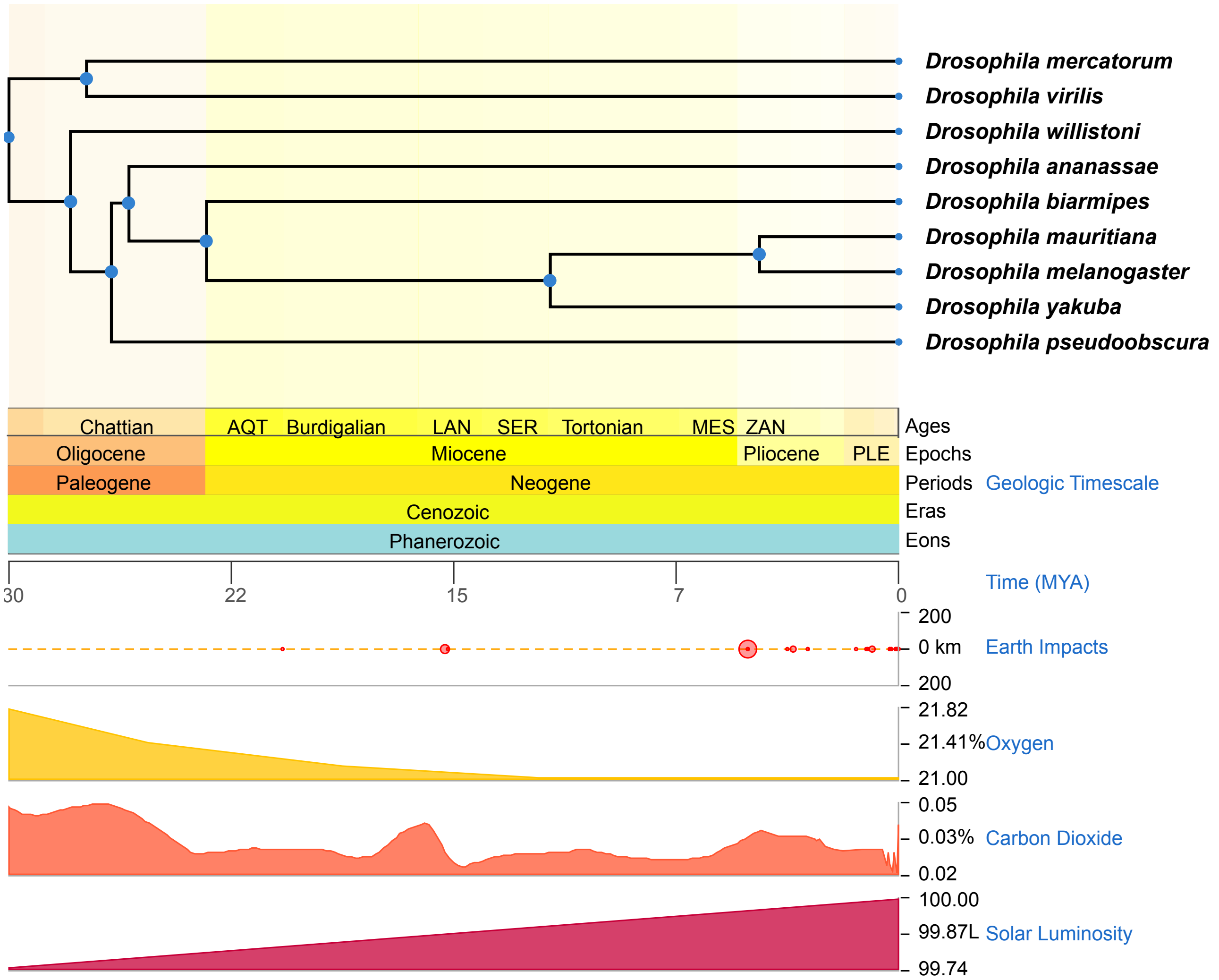


**Figure S1.** Phylogenetic tree of the species included in this study.

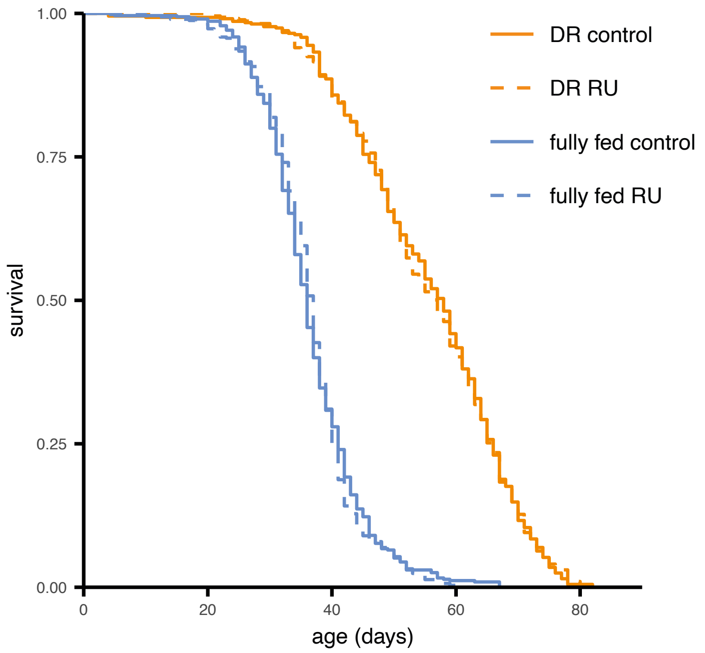


***Figure S1.*** Mifepristone (RU486) has no effect on survival on either fully fed (logHR = 0.053 ± 0.16, P = 0.74) or DR (logHR = -0.0035 ± 0.17, P = 0.98) conditions. Adult females tested using the exact same protocol as the longevity experiments using daughterless-GeneSwitch crossed to empty vector TRiP control line (attP2).
